# Cell-type specific transcriptomic signatures of neocortical circuit organization and their relevance to autism

**DOI:** 10.1101/2022.06.14.496156

**Authors:** Anthony J. Moussa, Jason C. Wester

## Abstract

A prevailing challenge in neuroscience is understanding how diverse neuronal cell types select their synaptic partners to form circuits. In the neocortex, major classes of excitatory projection neurons and inhibitory interneurons are conserved across functionally distinct regions. There is evidence these classes form canonical circuit motifs that depend primarily on their identity; however, regional cues likely also influence their choice of synaptic partners. We mined the Allen Institute’s single-cell RNA-sequencing database of mouse cortical neurons to study the expression of genes necessary for synaptic connectivity and maintenance in two regions: the anterior lateral motor cortex (ALM) and the primary visual cortex (VISp). We used the Allen’s metadata to parse cells by clusters representing major excitatory and inhibitory classes that are common to both ALM and VISp. We then performed two types of pairwise differential gene expression analysis: 1) between different neuronal classes within the same brain region (ALM or VISp), and 2) between the same neuronal class in ALM and VISp. We filtered our results for differentially expressed circuit connectivity related genes and developed a novel bioinformatic approach to determine the sets uniquely enriched in each neuronal class in ALM, VISp, or both. This analysis provides an organized set of genes that may regulate circuit formation in a cell-type-specific manner. Furthermore, it identifies candidate mechanisms for the formation of circuits that are conserved across functionally distinct cortical regions or that are region dependent. Finally, we used the SFARI Human Gene Module to identify genes from this analysis that are related to risk for autism spectrum disorder (ASD). Our analysis provides clear molecular targets for future studies to understand neocortical circuit organization and abnormalities that underly autistic phenotypes.

## INTRODUCTION

In the neocortex, excitatory projection neurons and inhibitory interneurons can be grouped into major classes that are found across functionally distinct regions. Excitatory classes are defined by their long-range axonal projection: intratelencephalic (IT), pyramidal tract (PT), or corticothalamic (CT) (Harris and Shepherd, 2015). Interneuron classes are defined by the expression of molecular markers such as parvalbumin (PV), somatostatin (SST), and vasoactive intestinal peptide (VIP) (Tremblay et al., 2016). Importantly, recent work suggests that neurons from these classes form microcircuit motifs that may be repeated across the cortex. For example, in prefrontal, motor, and visual cortices, IT cells from synaptic connections onto PT cells that are largely unreciprocated. (Morishima and Kawaguchi, 2006;Brown and Hestrin, 2009b;Kiritani et al., 2012). Among interneurons, VIP+ cells preferentially inhibit neighboring SST+ cells in primary sensory and prefrontal cortices (Lee et al., 2013;Pfeffer et al., 2013;Pi et al., 2013;Karnani et al., 2016). Furthermore, in deep cortical layers, PV+ interneurons preferentially target PT cells relative to IT cells (Lee et al., 2014;Wu et al., 2016;Ye et al., 2016), and VIP+ interneurons are targeted by IT cells but not PT cells (Wester et al., 2019). Thus, cell class appears to serve an important role in organizing canonical circuits for fundamental computations throughout the cortex (Douglas and Martin, 2004;Harris and Shepherd, 2015;Luo, 2021).

Recent single-cell RNA-sequencing studies have revolutionized our understanding of neocortical cell types and provide candidate molecular targets to study their synaptic connections (Zeisel et al., 2015;Tasic et al., 2016;Paul et al., 2017;Saunders et al., 2018;Tasic et al., 2018;Yao et al., 2021). These data reveal many subtypes of neurons within each major class, setting a foundation to investigate circuits with higher precision (Huang and Paul, 2019). However, they also raise questions regarding how to define the major classes in different cortical regions, which has important implications for understanding circuit organization. Specifically, excitatory classes vary in their transcriptomic profiles from the rostral to caudal poles (Saunders et al., 2018;Tasic et al., 2018;Yao et al., 2021). Thus, PT cells in primary visual cortex and prefrontal cortex may be best matched, but still categorized as different cell-types. In contrast, the transcriptomic profiles of interneuron classes are consistent throughout the cortex (Saunders et al., 2018;Tasic et al., 2018;Yao et al., 2021). This suggests that circuit motifs involving excitatory classes may be more regionally specialized than those involving interneurons. However, the extent to which the connectivity patterns of excitatory or inhibitory neurons can be generalized is unclear and an area of active research (Brown and Hestrin, 2009a;Huang and Paul, 2019;Luo, 2021). Indeed, recent work argues that circuits involving major interneuron classes are also dependent on region (Pouchelon et al., 2021). Thus, understanding how intrinsic class properties and areal cues guide the self-assembly of circuits remains a challenge.

Resolving these issues is necessary to design targeted approaches to treat neuropsychiatric disorders. For example, autism spectrum disorder (ASD) is hypothesized to result from an imbalance in the ratio of excitation to inhibition (E/I) within cortical circuits (Sohal and Rubenstein, 2019). Importantly, impairments that characterize ASD range from aberrant social behavior (entailing prefrontal circuits) (Yizhar et al., 2011) to sensory processing deficits (entailing visual cortical circuits) (Del Rosario et al., 2021). Thus, a tantalizing hypothesis is that in some instances ASD results from disruption of a canonical circuit motif that is repeated across functionally distinct regions. Alternatively, if circuit motifs in each region are unique, this would complicate therapeutic strategies. Several groups are now using monogenetic mouse models to investigate the contributions of different classes of excitatory and inhibitory neurons to the emergence of ASD phenotypes (Brumback et al., 2018;Mossner et al., 2020;Smith et al., 2020). However, clinically relevant mechanisms for synaptic connectivity, and if they are region specific, are unclear.

Here, we analyzed single-cell RNA-sequencing data from the Allen Institute for Brain Science (Tasic et al., 2018) to investigate molecular signatures of circuit organization in major neuronal classes across the neocortex. We used data from two cortical regions at opposite ends of the rostral-caudal axis to compare extremes in expression profiles: the motor planning anterior lateral motor cortex (ALM) and sensory processing primary visual cortex (VISp). Our analysis identifies synaptic genes that are enriched in select classes of excitatory and inhibitory cells. Furthermore, we determine which of these patterns are conserved in ALM and VISp, or unique to each region. Finally, we highlight classes that likely harbor specific ASD risk genes across the cortex and thus may be candidates for understanding constellations of conditions.

## RESULTS

### Transcriptomic profiles of both excitatory and inhibitory neurons are primarily differentiated by class rather than brain region

To compare neuronal subtypes between two functionally distinct cortical regions, we downloaded the Allen Institute’s single-cell RNA-sequencing datasets of adult mouse cortical neurons collected from the anterior lateral motor cortex (ALM) and primary visual cortex (VISp) (Tasic et al., 2018) (**Fig. 1A**). These include metadata that define each cell’s brain region (ALM or VISp), neurotransmitter (glutamatergic or GABAergic), and major class (e.g., L2/3 IT for excitatory neurons or Pvalb for inhibitory interneurons). We loaded these data into the R toolkit Seurat (Stuart et al., 2019) and ran the non-linear dimensionality reduction algorithm UMAP for unbiased clustering of excitatory projection neurons and inhibitory interneurons (**Figs. 1B, C**). We used the metadata to label each cell according to its brain region (**Figs. 1Bi, Ci**) or major class (**Figs. 1Bii,Cii**). As expected, this reproduced the findings of Tasic et al. (2018) that excitatory neurons cluster according to both brain region and class (**Fig. 1B**) but interneurons cluster only by class (**Fig. 1C**). This suggests that major classes of excitatory neurons (e.g., L2/3 IT) are not conserved across cortical regions but are distinct in ALM and VISp (Tasic et al., 2018). However, relative positions of clusters in UMAP space do not indicate the magnitude of differences between cell-types. Thus, we performed differential gene expression analyses to compare the number of genes that distinguish neuronal classes within and across brain regions (**Figs. 2A, B**).

**Figure 1.**
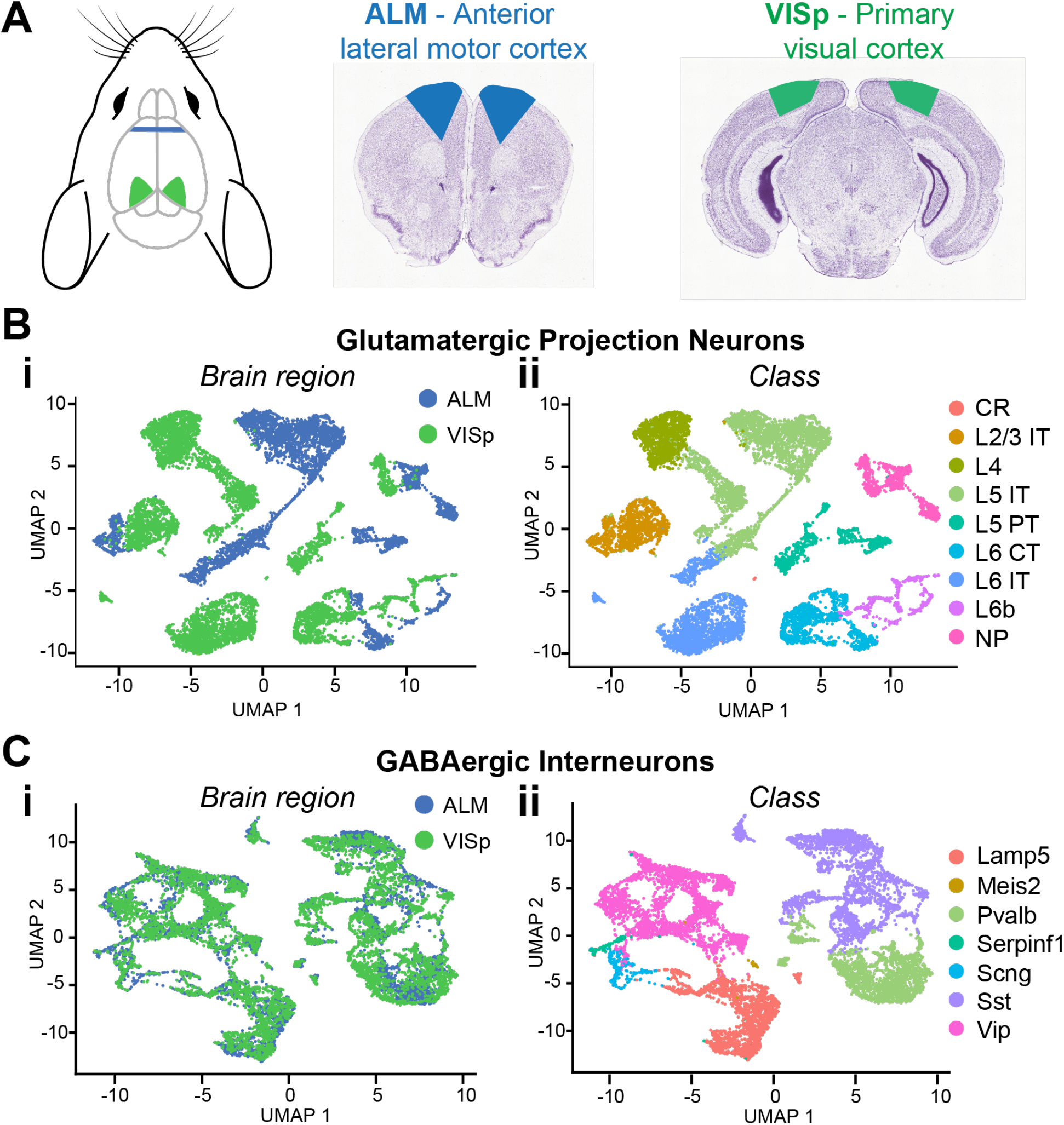
Single-cell RNA-seq data for major neuronal classes in ALM and VISp. **(A)** Anatomical locations of ALM (blue) and VISp (green) with corresponding coronal sections. **(B)** UMAP dimensionality reduction for glutamatergic cells color-coded using metadata for brain region **(Bi)** or class **(Bii). (C)** UMAP dimensionality reduction for GABAergic cells color-coded using metadata for brain region **(Ci)** or class **(Cii)**. L2/3 IT, L5 IT, L5 PT, L6 CT, Pvalb, Sst, and Vip clusters were selected for downstream analysis. CR, Cajal–Retzius; IT, intratelencephalic; PT, pyramidal tract; CT, corticothalamic; NP, near-projecting; Pvalb, parvalbumin; Sst, somatostatin; Vip, vasoactive intestinal peptide. Coronal sections adapted from the Allen Mouse Brain Atlas (https://mouse.brain-map.org/static/atlas).

**Figure 2.**
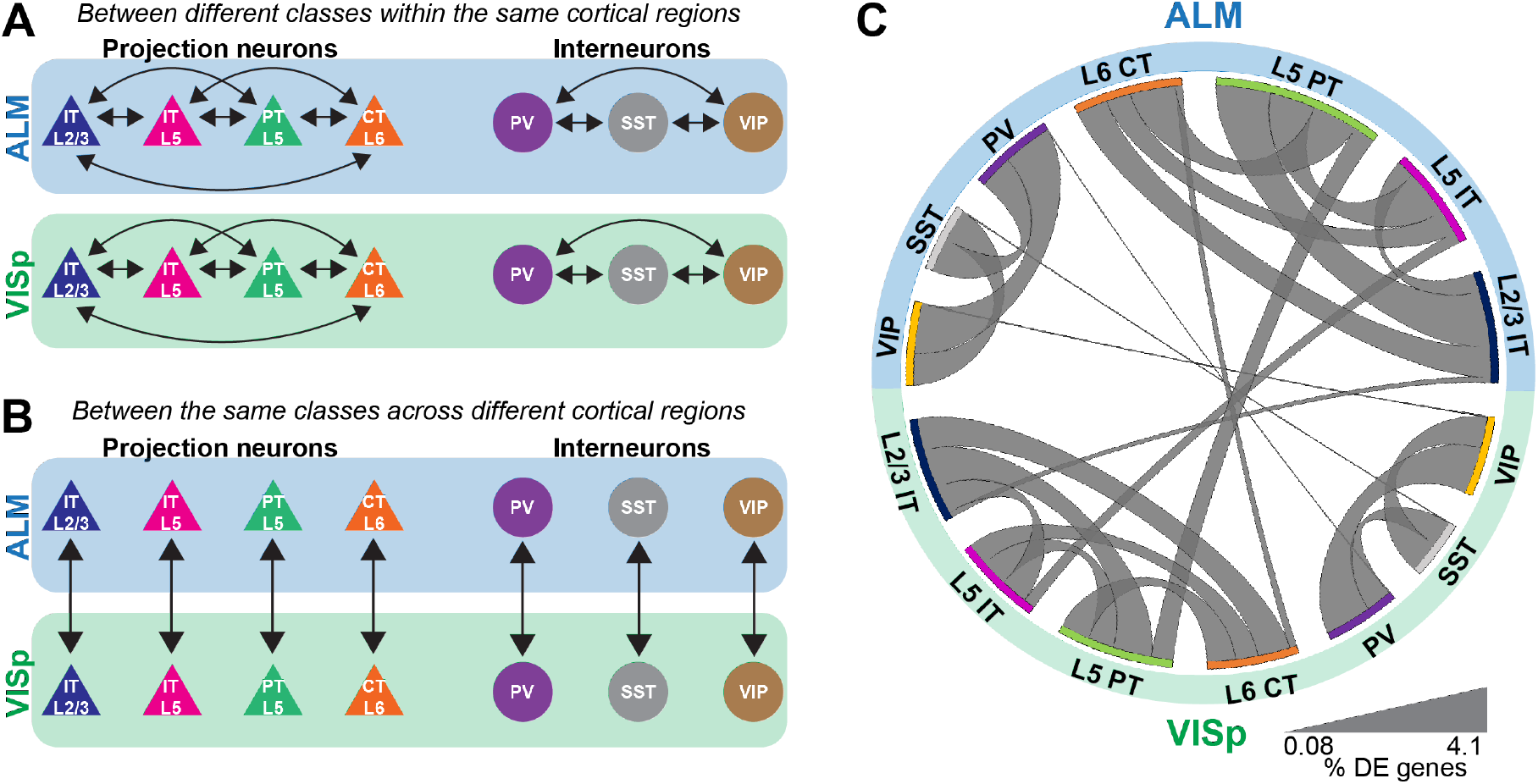
Differential expression tests for major neuronal classes either within or between cortical regions. **(A)** Pairwise differential expression tests conducted between excitatory or inhibitory neuronal classes in ALM or VISp. **(B)** Pairwise differential expression tests conducted between the same neuronal classes across ALM or VISp. **(C)** Circos plot summary of all differential expression test results. These include results for all transcripts in the dataset, in contrast to Figure 3 in which genes are filtered by gene ontology labels. Classes are listed along the circumference of the plot in their respective brain region (blue - ALM; green - VISp). Bands connect classes that were compared in a pairwise differential expression test. Band thickness represents the proportion of differentially expressed genes in each test: thinner bands indicate a small percentage of distinguishing genes and thicker bands indicate a larger percentage of distinguishing genes.

For our analyses, we chose a subset of major excitatory and inhibitory neuronal classes that are shared between ALM and VISp and are relatively well studied (Harris and Shepherd, 2015;Tremblay et al., 2016). For excitatory cells, these included layer 2/3 intratelencephalic (L2/3 IT), layer 5 intratelencephalic (L5 IT), layer 5 pyramidal tract (L5 PT), and layer 6 corticothalamic (L6 CT). For inhibitory cells, these included those expressing parvalbumin (PV), somatostatin (SST), or vasoactive intestinal peptide (VIP). We conducted two types of analyses: 1) pairwise differential expression tests between different excitatory or inhibitory classes within the same cortical region (ALM or VISp) (**Fig. 2A**) and 2) pairwise differential expression tests between the same classes across ALM and VISp (**Fig. 2B**). For each pairwise test, we counted the number of differentially expressed genes and normalized it to the total number compared (**Supplemental Spreadsheet 1, “Circos Table” “All Genes” columns)**. This allowed us to determine the proportions of differentially expressed genes among all of the pairwise tests both within and across brain regions, which we visualized using a Circos plot (Krzywinski et al., 2009) (**Fig. 2C**). For both excitatory and inhibitory cells, we found that each major class is more similar between brain regions than between other classes within the same brain region. Thus, even though excitatory neurons cluster by brain region (**Fig. 1B**), class identity best distinguishes them. Interestingly, we also found differences between major inhibitory classes across ALM and VISp, suggesting that regional cues also influence their differentiation (Pouchelon et al., 2021). Our results agree with previous work that major excitatory and inhibitory classes are primarily conserved across diverse cortical regions when considering transcriptomic profiles (Tasic et al., 2018;Yao et al., 2021).

### Genes relevant to synaptic connectivity are primarily differentiated by class rather than brain region

Although major neuronal classes largely share transcriptomic profiles between ALM and VISp, these two cortical regions are functionally distinct. Thus, we next asked if genes related to synaptic connectivity and circuit organization are also shared for each neuronal class across the cortex. We filtered our differential expression results using the gene ontology (GO) software PANTHER 16.0 (Mi et al., 2021) to select genes labeled by the terms “Cell-cell adhesion”, “Regulation of cell-cell adhesion”, and “Regulation of trans-synaptic signaling”. These were chosen because they include genes related to the formation, maintenance, and modulation of synaptic connections among distinct neuronal classes (Fuccillo et al., 2015;Földy et al., 2016;Chowdhury et al., 2021). Throughout, we refer to these as “circuit-related” genes. For all three GO terms, we found lower proportions of differentially expressed circuit-related genes between the same class in ALM and VISp than between different classes, regardless of brain region **(Fig. 3; Supplemental Spreadsheet 1, “Circos Table”)**. This matches the pattern observed for differential expression of all transcripts (**Fig. 2C**). Thus, these analyses suggest that despite the functional distinction of ALM and VISp, gene expression profiles that define cell-types and their markers for circuit organization are largely conserved across the neocortex.

**Figure 3.**
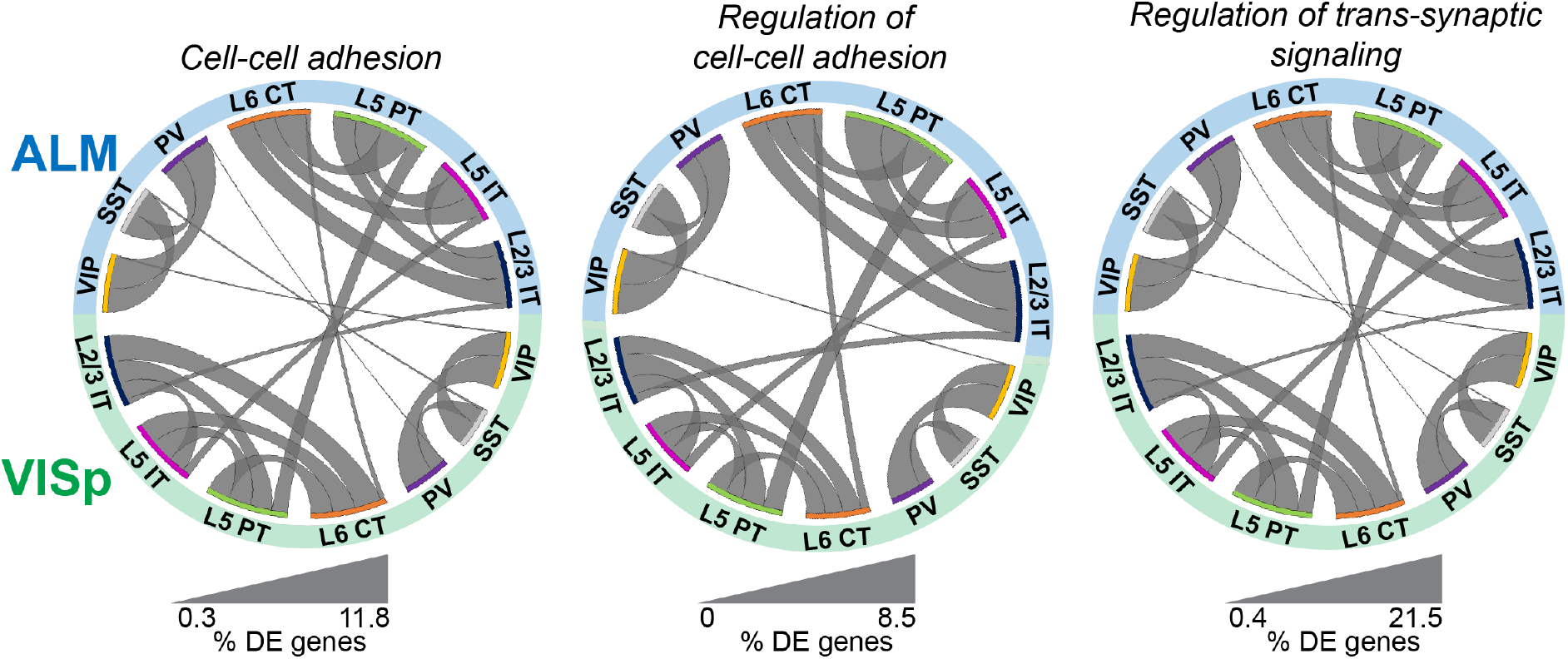
Circuit-related genes are primarily differentiated by class rather than cortical region. Circos plots of differential expression results filtered for circuit-related genes using PANTHER gene ontology labels “cell-cell adhesion”, “regulation of cell-cell adhesion”, and “regulation of trans-synaptic signaling”. Classes are listed along the circumference of the plot in their respective brain region (blue - ALM; green - VISp). Bands connect classes that were compared in a pairwise differential expression test. Band thickness represents the proportion of differentially expressed genes in each test: thinner bands indicate a small percentage of distinguishing genes and thicker bands indicate a larger percentage of distinguishing genes. No genes involved in Regulation of cell-cell adhesion were differentially expressed between SST or PV cells across regions.

### Identifying the circuit-related genes that are differentially expressed between classes reveals that most are region-specific for excitatory projection neurons

The above analyses simply consider the proportions of differentially expressed genes to describe general trends in the data. Thus, by percentage most circuit-related genes are differentially expressed between neuronal classes within ALM and VISp rather than between the same classes across these cortical regions (**Fig. 3**). However, this analysis does not consider if the genes that are differentially expressed between neuronal classes within ALM and VISp are different sets. This could obscure the true number of genes that provide region-specific instructions for circuit organization. Thus, for each differential expression test between neuronal classes within ALM or VISp, we categorized the result as either 1) ALM-specific, 2) VISp-specific, 3) conserved, or 4) unique (**Fig. 4A**). For example, *rock1* is differentially expressed between VIP and PV cells in ALM but not VISp (ALM-specific, **Fig. 4Ai**). Similarly, *reln* is differentially expressed between VIP and PV cells in VISp but not ALM (VISp-specific, **Fig. 4Aii**). Importantly, for a gene that is differentially expressed between the same pair of neuronal classes in both ALM and VISp, it is necessary to consider which class had greater expression in each comparison. For example, *shisa6* exhibits greater expression in VIP cells relative to PV cells in both ALM and VISp (conserved, **Fig. 4Aiii**). However, *calb1* exhibits greater expression in L5 IT cells relative to L5 PT cells in ALM but this relationship is reversed in VISp (unique, **Fig. 4Aiv**).

**Figure 4.**
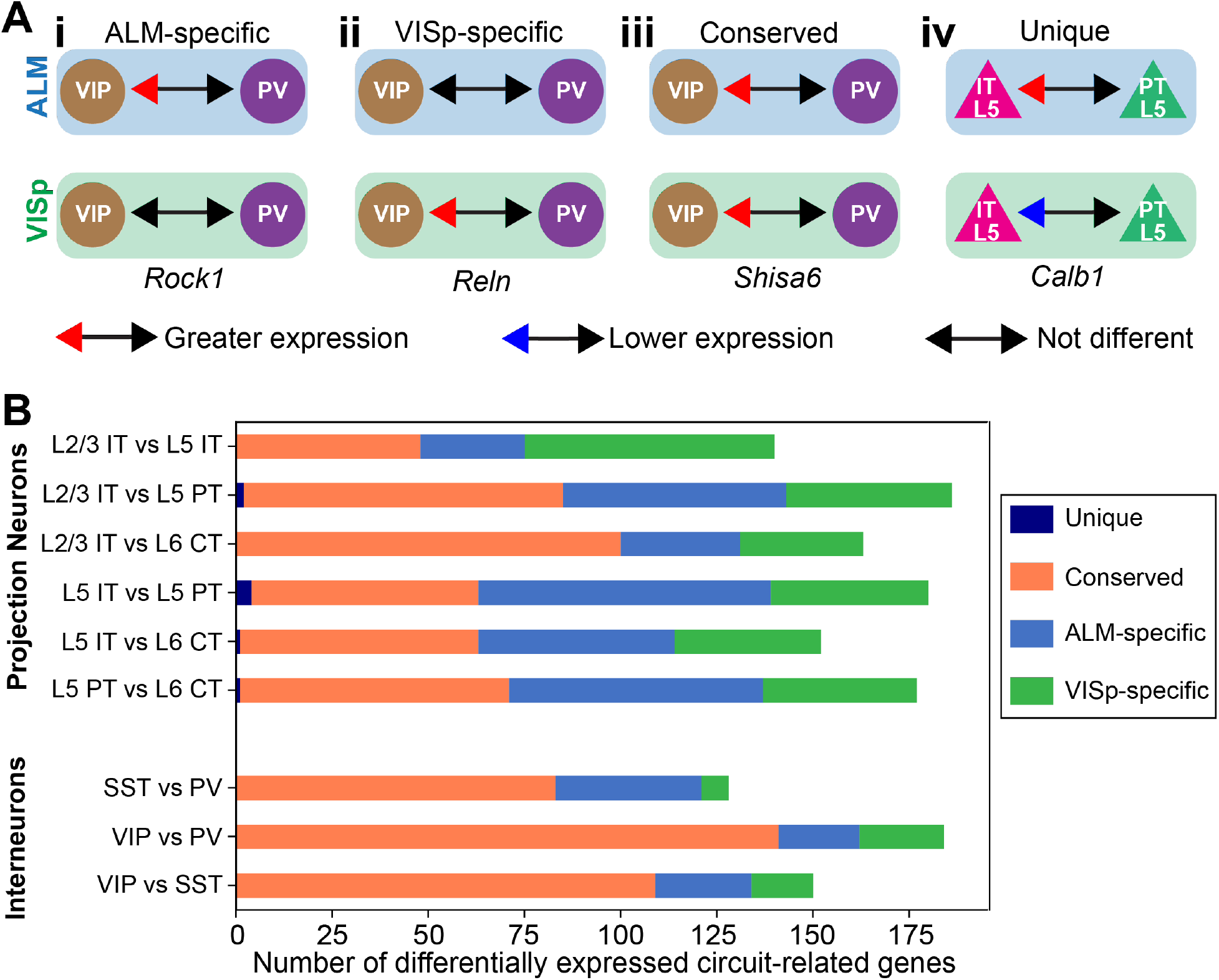
Categorization of circuit-related genes that are differentially expressed between neuronal classes within ALM or VISp. **(A)** Examples of differential expression test results used to categorize circuit-related genes as: ALM-specific **(Ai)**, VISp-specific **(Aii)**, Conserved **(Aiii)**, or Unique **(Aiv)**. The genes and results are real examples from the analysis. **(B)** Stacked bar plot of the total number of circuit-related genes in each category for every pairwise differential expression test between classes. “Unique” genes were only observed in L2/3 IT vs L5 PT; L5 IT vs L5 PT; L5 IT vs L6 CT; and L5 PT vs L6 CT comparisons.

In **Figure 4B**, we plotted the total number of circuit-related genes in each category that demonstrated significant differential expression between neuronal classes. Importantly, genes in the “unique” category were rare and only observed in a subset of tests between excitatory projection neurons (only 7 genes in 4 tests). Thus, the analysis from our differential expression tests yielded few results that would require more complicated interpretations by considering each brain region separately. As expected, for inhibitory interneurons most differential expression test results were conserved between ALM and VISp. This is consistent with the observation that interneuron expression profiles do not cluster by brain region (**Fig. 1C**) and our Circos plots of proportions of differentially expressed genes within and across cortical regions (**Figs. 2C, 3**). However, we identified several genes for which differential expression results were ALM- or VISp-specific and consider them in detail in below (**Figs. 8 and 9**). Strikingly, for excitatory neurons we found that more than half of the circuit-related genes in each comparison were exclusive to ALM or VISp (except for the L2/3 IT vs L6 CT comparison). This appears incongruent with our above results that excitatory classes have a lower proportion of differentially expressed circuit-related genes across brain regions (**Fig. 3**). However, we emphasize that this analysis is unique from **Figures 2 and 3**. In **Figure 4**, we consider the *identity* of circuit-related genes that are differentially expressed between neuronal classes *within* each brain region (as in **Fig. 2A**) to determine if the gene sets and their differential expression results overlap. This reveals that for excitatory projection neurons most circuit-related genes that distinguish classes within ALM and VISp are exclusive to each region.

### The results from individual differential expression tests can be combined to identify circuit-related genes relevant to each major neuronal class

In the above analysis, we found 1452 significant differential expression test results for 394 circuit-related genes that were exclusive to ALM, exclusive to VISp, or conserved between cortical regions (**Fig. 4B; Supplemental Spreadsheet 1 “StackedBar Table”**). This makes it challenging to derive practical biological insights from these data. Thus, we developed a novel bioinformatic approach to narrow our results to a tractable set of genes that have a high likelihood to play key roles in cortical circuit organization. Our strategy identified circuit-related genes for which differential expression results were consistent across all tests for each neuronal class. As an example, we consider a subset of seven circuit-related genes with significant differential expression among inhibitory neuronal classes in **Figure 5**. *Cyfip1* has greater expression in VIP cells than PV cells in ALM, but not in VISp (**Fig. 5A**); thus, we classified this gene as ALM-specific (**Fig. 5Bi**). *Dlg2* has greater expression in VIP cells than PV cells in *both* ALM and VISp (**Fig. 5A**); thus, we classified this gene as Conserved (**Fig. 5Bii**). Finally, *kctd13* has greater expression in VIP cells than PV cells in VISp, but not in ALM (**Fig. 5A**); thus, we classified this gene as VISp-specific (**Fig. 5Biii**). For these three genes, differential expression between SST and PV cells is not significantly different; thus, we would predict greater expression in VIP cells relative to SST cells for each. However, this was not the case (**Fig. 5A**), which renders the biological relevance of these results unclear. In contrast, *nrxn1, pten, snap25*, and *socs5*, all demonstrate consistent differential expression results among all three tests (**Fig. 5A**). Thus, we sorted these genes as ALM-specific (*socs5*), VISp-specific (*pten*), or Conserved (*nrxn1* and *snap25*), as above (**Fig. 5B**). We then kept these four genes for further analysis, and discarded *cyfip1, dlg2*, and *kctd13* (**Fig. 5C**). Next, we defined *nrxn1, pten, snap25*, and *socs5* as “class-relevant” to VIP cells and denoted whether expression was higher or lower relative to the other classes with up and down arrows, respectively (see **Fig. 5Cii**). Finally, dysregulation of circuit-related genes directly contributes to neurological disorders (Chowdhury et al., 2021). Thus, we highlighted in green candidate risk genes for autism using the SFARI Human Gene Module (https://gene.sfari.org/database/human-gene/) (**Fig. 5C**).

**Figure 5.**
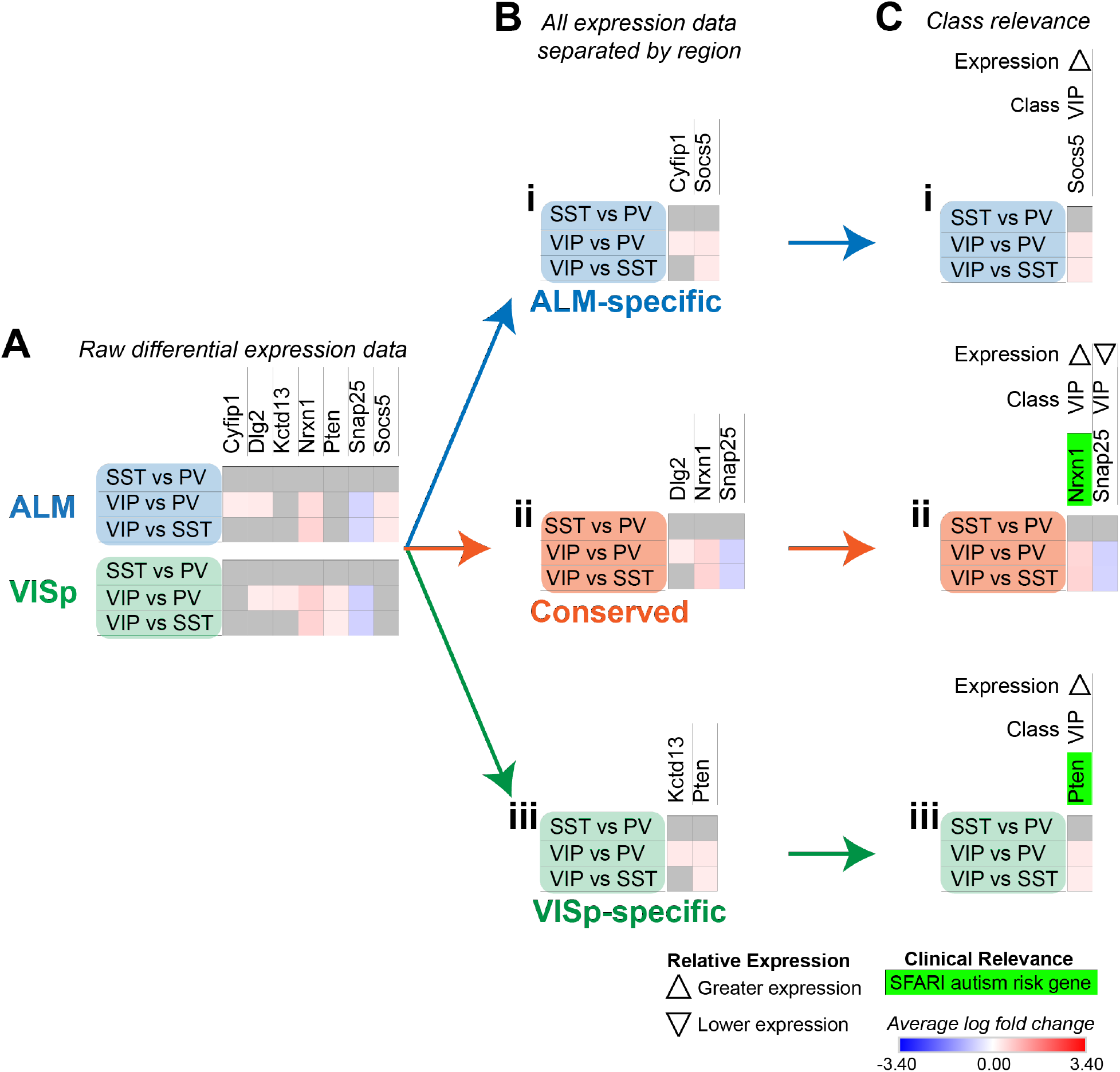
Strategy to determine circuit-related genes relevant to specific neuronal classes. **(A)** Examples of differential expression rests results for seven circuit-related genes in interneuron classes in ALM and VISp. **(B)** Separation of genes from *(A)* into ALM-specific **(Bi)**, Conserved **(Bii)**, and VISp-specific **(Biii)** subsets. **(C)** Class-relevant genes from *(B)* identified by their consistent differential expression results among all class tests. For each gene, we provide a class-relevant label and an arrow indication whether it is expressed higher or lower relative to other classes. Class-relevant genes listed on the SFARI Human Gene Module as implicated in autism were highlighted green.

For some class-relevant genes, a subset of their differential expression results was shared between ALM and VISp, and a subset was exclusive to one cortical region. As an example, we include the analysis of *egr1* in **Figure 6**. *Erg1* demonstrates significant differential expression in all three tests between interneuron classes in ALM, but in only two tests in VISp (**Fig. 6A**). Thus, in ALM *erg1* has greater expression in VIP cells relative to SST cells (**Fig. 6Bi**), while in both ALM and VISp *erg1* has greater expression in SST cells relative to PVs cells and in VIP cells relative to PV cells (**Fig. 6Bii**). These differences are important for interpreting the relevance of *erg1* expression among neuronal classes. In ALM, we define *erg1* as having greatest expression in VIP cells relative to PV and SST cells (**Fig. 6Ci**). However, we indicate with an asterisk (*) that a subset of these results is conserved in VISp, where we define *erg1* as having lower expression in PV cells relative to VIP and SST cells (**Fig. 6Cii**).

**Figure 6.**
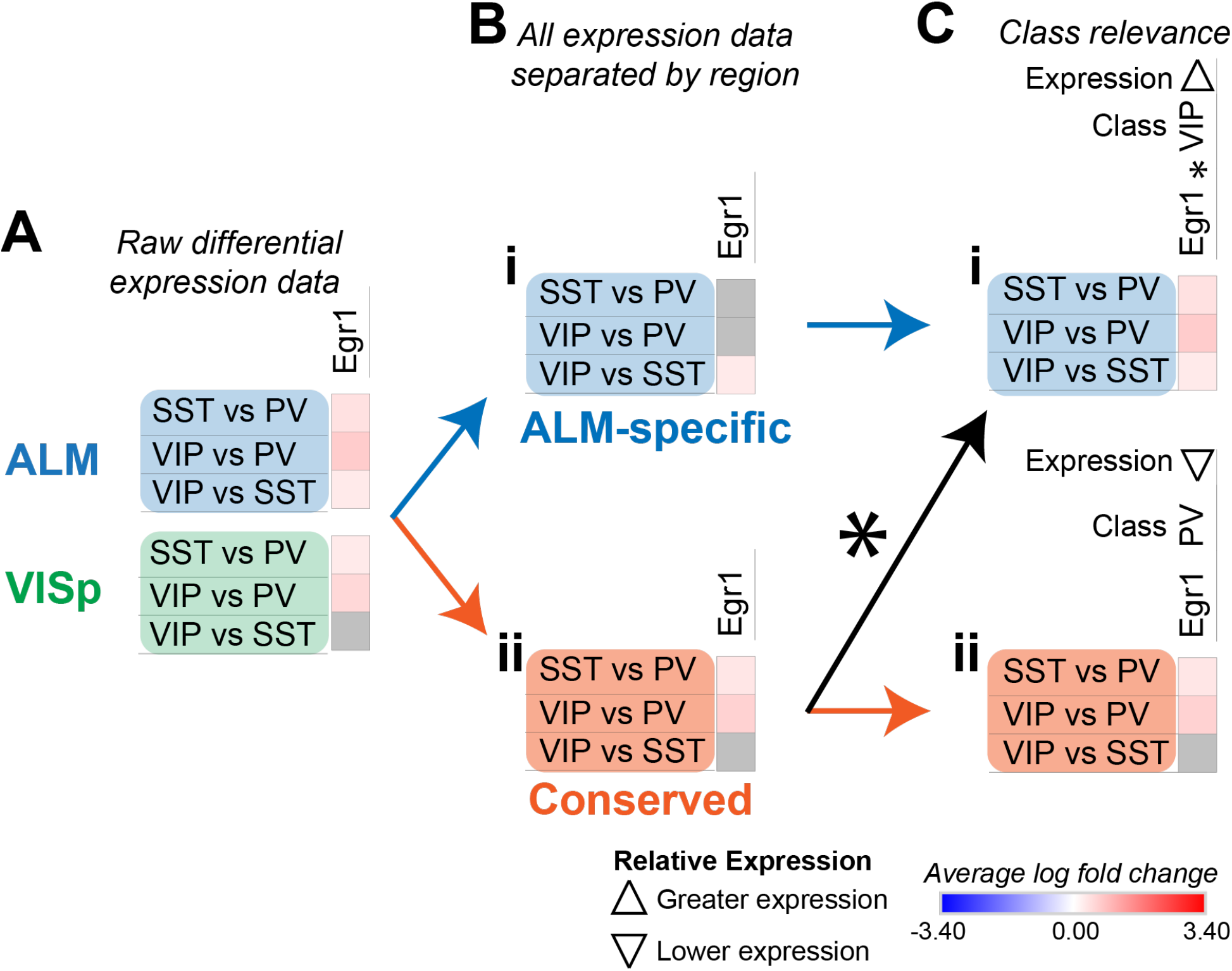
Strategy to identify class-relevant genes that describe both conserved and region-specific relationships. **(A)** Example differential expression test results for *egr1* in both ALM and VISp. **(B)** Separation of differential expression results into subsets that are ALM-specific **(Bi)** and conserved in both ALM and VISp **(Bii). (C)** Annotation of *erg1* using an asterisk (*) to denote that a subset of ALM-specific differential expression results **(Ci)** is also found within the set labeled as “Conserved” **(Cii)**. Note that for *erg1*, the relevant class and relative expression-level are different in *Ci* and *Cii* because the combinations of differential expression results among neuronal classes are unique.

In summary, our strategy combined pairwise differential expression test results to relate each circuit-related gene to a major neuronal class. This allowed us to create an organized and tractable list of genes that are candidates to mediate synaptic connections between specific neuronal classes across the cortex or dependent on cortical region (**Figs. 7 – 9**). Of the 394 circuit-related genes in the ALM-specific, VISp-specific, and Conserved subsets, 71% (280) mapped to at least one class. Furthermore, 26% of these are risk genes for autism and may inform future studies investigating the circuit mechanisms of this disorder. We describe these data in greater detail below and provide a searchable spreadsheet that identifies each gene-class relationship (**Supplemental Spreadsheet 2**).

**Figure 7.**
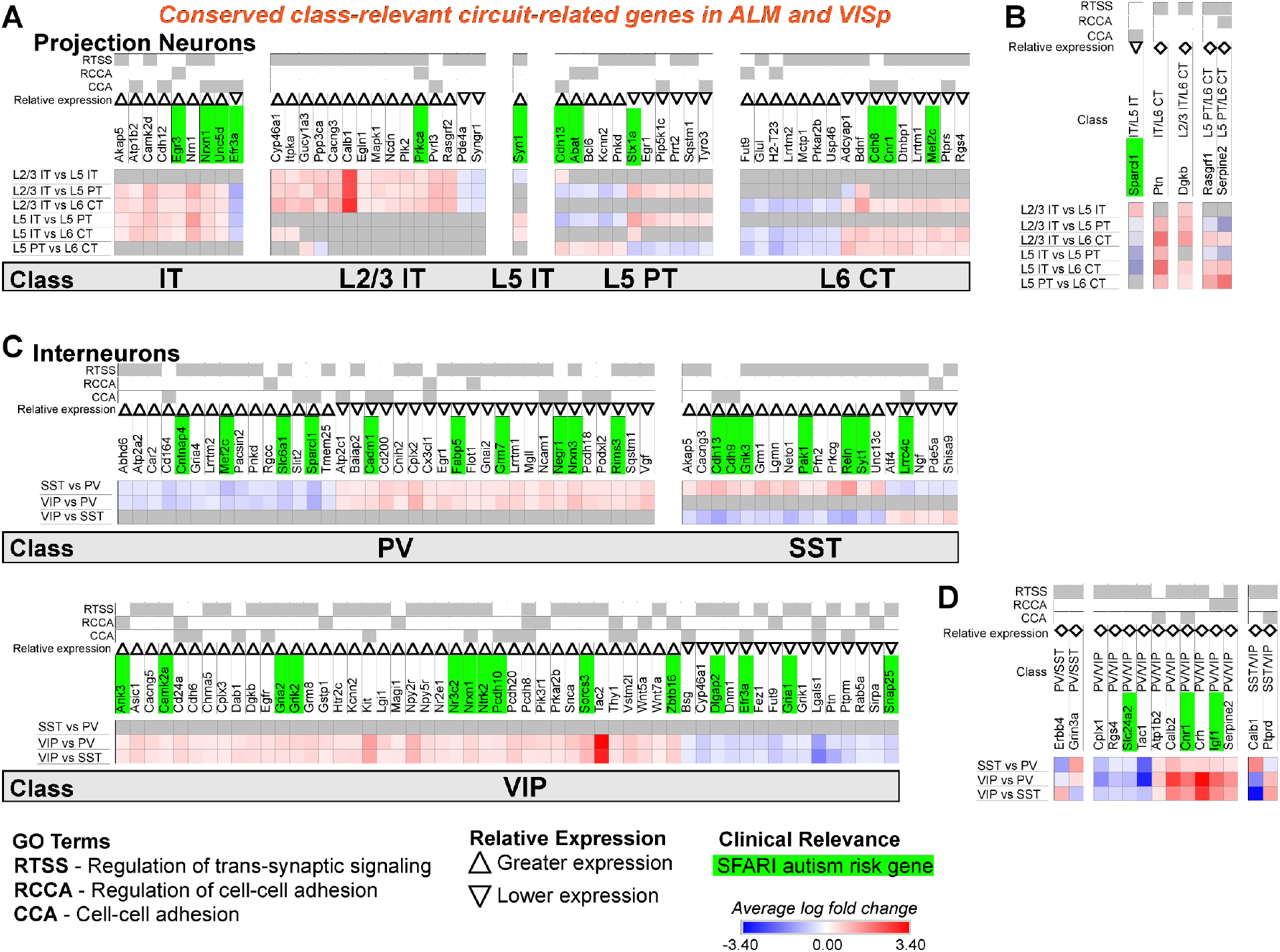
Circuit-related genes that are relevant to the same neuronal classes in ALM and VISp. **(A)** Circuit-related genes relevant to single classes of excitatory projection neurons. We created the new class “IT” for genes that were relevant to both L2/3 IT and L5 IT classes. **(B)** Circuit-related genes relevant to multiple classes of excitatory projection neurons. **(C)** Circuit-related genes relevant to single classes of inhibitory interneurons. **(D)** Circuit-related genes relevant to multiple classes of inhibitory interneurons. CCA = cell-cell adhesion; RCCA = regulation of cell-cell adhesion; RTSS = regulation of trans-synaptic signaling.

### Most circuit-related genes identified as Conserved are class-relevant to inhibitory interneurons

Our analysis identified 57 circuit-related genes relevant to excitatory neuronal classes that were conserved in both ALM and VISp (**Fig. 7A, B**). A subset of 9 genes was relevant to both L2/3 IT and L5 IT classes; thus, we created a new class “IT” for these genes (**Fig. 7A**). Indeed, most circuit-related genes that were conserved among excitatory neurons in both ALM and VISp were relevant to the L2/3 IT, L5 IT, and combined IT classes. Only a small number of genes (5) were relevant to other combinations of multiple classes (e.g., L2/3 IT and L6 CT) in both ALM and VISp (**Fig. 7B**). Interestingly, more than twice as many genes (124) were conserved for interneuron classes compared to excitatory classes (**Fig. 7C, D**). Furthermore, most of these genes were unique to interneurons; only 23 genes were identified as class-relevant in both the excitatory and inhibitory populations. Thus, the mechanisms that regulate synaptic connections in a class-specific manner may be mostly distinct between excitatory and inhibitory cell types. Among interneurons, most were relevant to VIP cells and distinguished them from PV and SST cells (**Fig. 7C**). Thus, the mechanisms underlying the formation of disinhibitory circuits appear to be conserved across cortical regions. Finally, SFARI autism risk genes were distributed uniformly among all excitatory and inhibitory classes and represented 23% and 28% of the total genes for each group, respectively. Thus, we could not identify a specific set of neuronal classes that may be uniquely vulnerable in autism. In summary, we identified circuit-related genes that are conserved between ALM and VISp for each neuronal class and may be molecular mechanisms underlying stereotyped “canonical circuits” found across the cortex (Kubota, 2014;Harris and Shepherd, 2015;Gutman-Wei and Brown, 2021). Furthermore, many of them are candidates to disrupt the balance between excitation and inhibition throughout the cortex in developmental disorders (Sohal and Rubenstein, 2019).

### Most circuit-related genes that are identified as specific to classes in ALM or VISp are in excitatory projection neurons

We next evaluated circuit-related genes that are relevant to neuronal classes exclusively in ALM (**Fig. 8**) or VISp (**Fig. 9**). In ALM, we found 92 genes relevant to excitatory classes (**Fig. 8A**), 15 of which shared a subset of differential expression results that were also relevant to classes in VISp (**Fig. 8A**, genes denoted by * are also found in **Fig. 7A**). Only a small number of genes (6) were relevant to combinations of multiple classes (e.g., L5 IT and L5 PT) exclusively in ALM (**Fig. 8B**). For interneurons, we found 47 class-relevant genes (**Fig. 8C**), 11 of which shared a subset of differential expression results that were also relevant to classes in VISp (**Fig. 8C**, genes denoted by * are also found in **Fig. 7C**). Like our analyses of conserved genes (**Fig. 7**), few were shared between excitatory and inhibitory classes (12 total). However, in contrast to conserved genes, most were class relevant to excitatory cells. Furthermore, among excitatory cells most genes were relevant specifically to the IT classes. Our analysis of genes that were class relevant in VISp yielded similar results (**Fig. 9**). In VISp, we found 79 genes relevant to excitatory classes (**Fig. 9A**), 12 of which shared a subset of differential expression results that were also relevant to classes in ALM (**Fig. 9A**, genes denoted by * are also found in **Fig. 7A**). Only 2 genes were relevant to combinations of multiple classes (e.g., L5 PT and L6 CT) exclusively in VISp (**Fig. 9B**). For interneurons, we found 28 class-relevant genes (**Fig. 9C**), 8 of which shared a subset of differential expression results that were also relevant to classes in ALM (**Fig. 9C**, genes denoted by * are also found in **Fig. 7C**). Like in ALM, most genes in VISp were relevant to excitatory IT-type classes, and few overlapped between excitatory and inhibitory cells (9 total).

**Figure 8.**
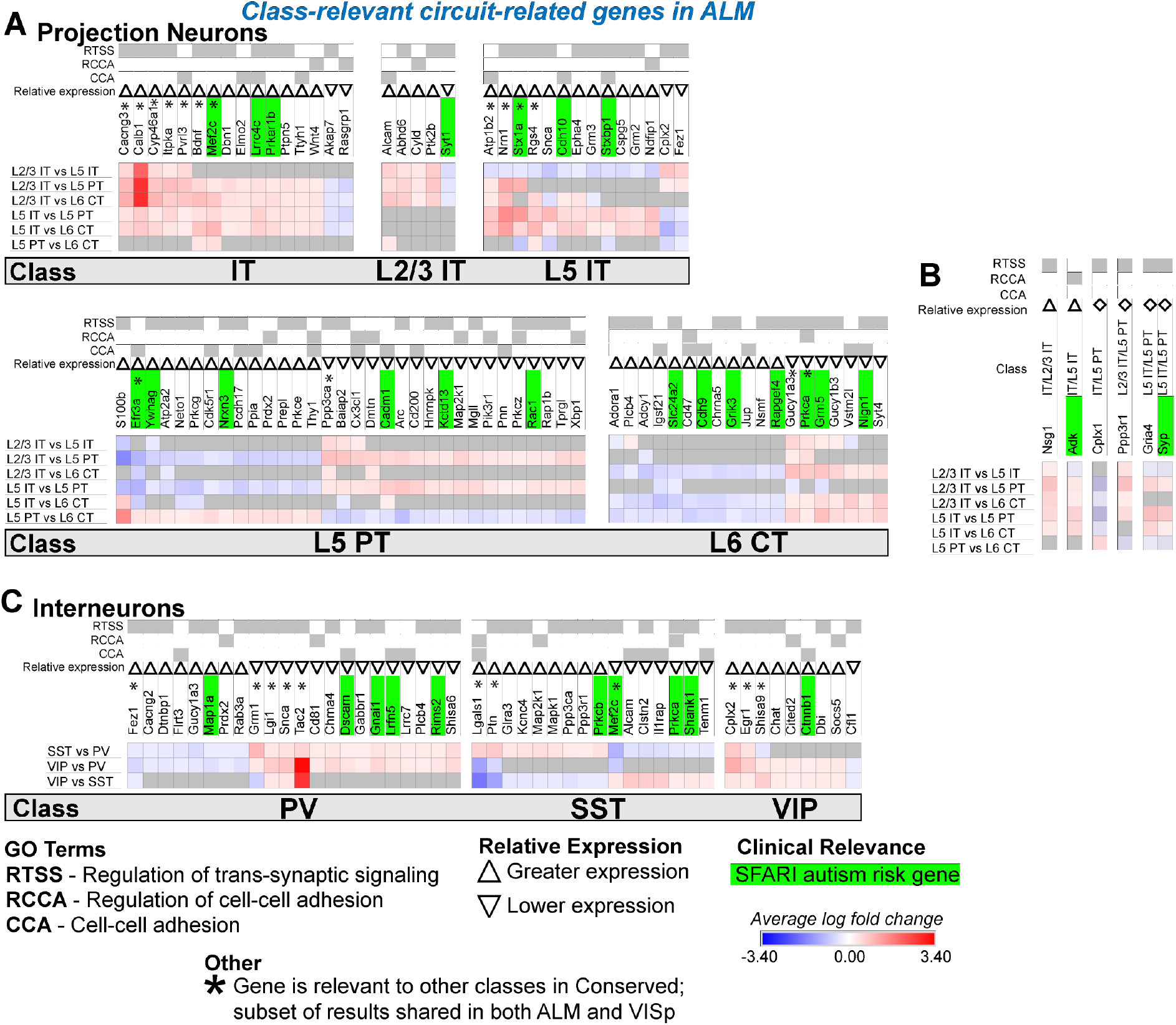
Circuit-related genes only relevant to neuronal classes in ALM. **(A)** Circuit-related genes relevant to single classes of excitatory projection neurons. We created the new class “IT” for genes that were relevant to both L2/3 IT and L5 IT classes. **(B)** Circuit-related genes relevant to multiple classes of excitatory projection neurons. **(C)** Circuit-related genes relevant to single classes of inhibitory interneurons. CCA = cell-cell adhesion; RCCA = regulation of cell-cell adhesion; RTSS = regulation of trans-synaptic signaling.

**Figure 9.**
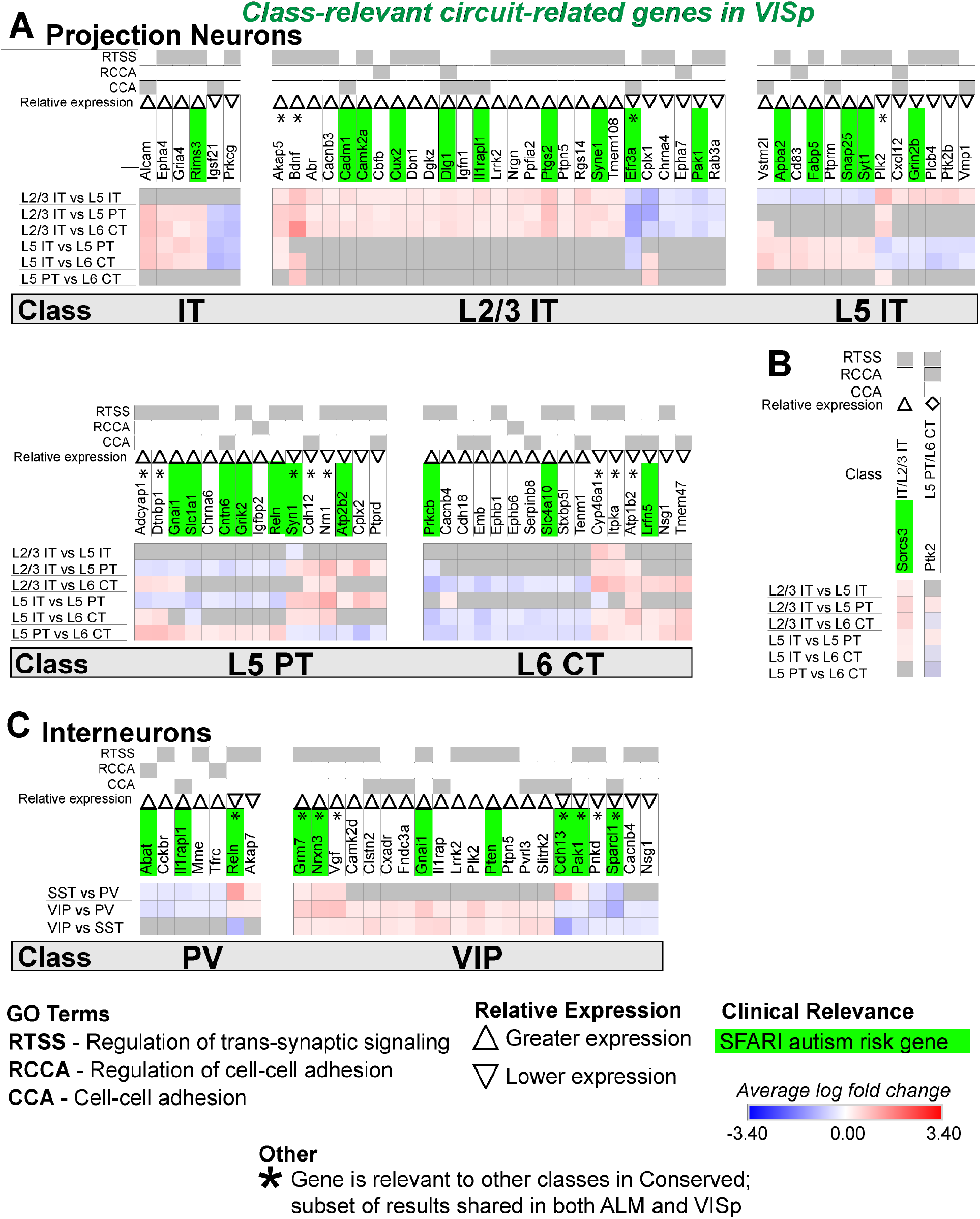
Circuit-related genes only relevant to neuronal classes in VISp. **(A)** Circuit-related genes relevant to single classes of excitatory projection neurons. We created the new class “IT” for genes that were relevant to both L2/3 IT and L5 IT classes. **(B)** Circuit-related genes relevant to multiple classes of excitatory projection neurons. **(C)** Circuit-related genes relevant to single classes of inhibitory interneurons. CCA = cell-cell adhesion; RCCA = regulation of cell-cell adhesion; RTSS = regulation of trans-synaptic signaling.

In summary, we found that most of the class-relevant circuit-related genes that are dependent on cortical region are found in excitatory neurons. Furthermore, most of these are relevant to IT-type cells and many are candidate risk genes for autism. These data agree with previous work in humans that IT-type cells may be a key contributor to autistic phenotypes (Parikshak et al., 2013). However, our results highlight that the development of therapies to target excitatory cells may not work uniformly across the cortex.

### A small set of genes demonstrate greater differential expression exclusively in ALM or VISp

Finally, we analyzed the differential expression results for each class between ALM and VISp (**Fig. 10**; see **Fig. 2B** for schematic). We hypothesized there are genes that are uniquely expressed between these functionally distinct and spatially distant cortical regions. Thus, we filtered genes that demonstrated consistent differential expression results for all neuronal classes between ALM and VISp. Strikingly, this analysis identified few genes. Among excitatory classes, only 4 genes were biased to ALM (*lmo4, lphn2, neurod6*, and *tmeff1*) and only 4 were biased to VISp (*brinp3, id2, spock3, tenm2*) (**Fig. 10A**). Only 1 of these genes, *tenm2*, was circuit-related (**Fig. 10A, D**). Among inhibitory classes, we found only 1 gene (*cenpa*) that was biased to VISp, but it was not circuit-related according to gene ontology (but see Discussion section) (**Fig. 10B, C**). Finally, PV and SST interneurons share their developmental origin from the medial ganglionic eminence (Xu et al., 2004;Butt et al., 2005), but VIP interneurons are derived from the caudal ganglionic eminence (Lee et al., 2010;Miyoshi et al., 2010). Thus, we asked if there are circuit-related genes that distinguish interneurons in ALM and VISp dependent on embryonic origin. We found none for PV/SST classes and only 4 such genes (*calb2, igf1, npy2r*, and *pvrl3*) for the VIP class (**Fig. 10B, E**). Of note, a previous study showed that *igf1* expression in VIP interneurons regulates experience-dependent plasticity in primary visual cortex (Mardinly et al., 2016). In summary, surprisingly few genes collectively distinguish excitatory and inhibitory neuronal classes between the rostral and caudal poles of the cortex.

**Figure 10.**
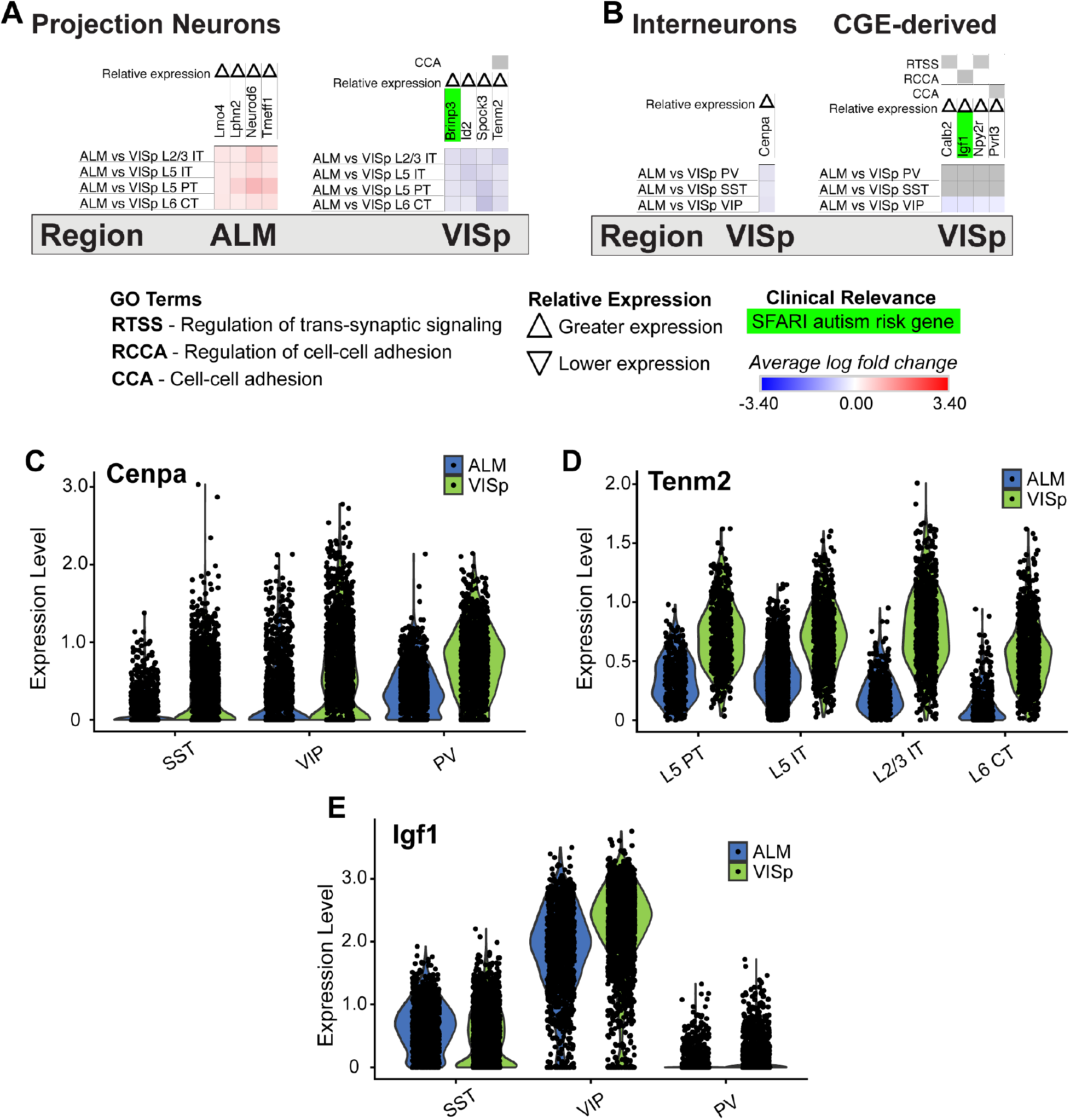
Genes that have biased expression to ALM or VISp for all excitatory or inhibitory neuronal classes. **(A)** Genes with biased expression between ALM or VISp for excitatory projection neuron classes. Note that only 1 gene, *tenm2*, is circuit-related. **(B)** Genes with biased expression between ALM or VISp for inhibitory interneuron classes. (*Left*) Only 1 gene, *cenpa*, had biased expression among all classes. (*Right*) Only 4 circuit-related genes demonstrated biases in expression when comparing VIP to PV/SST classes. **(C)** Violin plots of *cenpa* single-cell expression for inhibitory interneuron classes in ALM and VISp. **(D)** Violin plots of *tenm2* single-cell expression for excitatory neuronal classes in ALM and VISp. **(E)** Violin plots of *igf1* single-cell expression for inhibitory interneuron classes in ALM and VISp.

## DISCUSSION

### Regional differentiation of neuronal classes and implications for circuit organization

Excitatory projection neuron progenitors in the developing dorsal telencephalon are organized as a protomap of cortical regions (Rakic, 1988;Elsen et al., 2013;Greig et al., 2013). This protomap includes gradients of transcription factors (O’Leary et al., 2007;Cadwell et al., 2019) that vary from the rostral to caudal poles and may give rise to distinct lineages of excitatory neurons (Nowakowski et al., 2017;Bhaduri et al., 2021). Thus, it is not surprising that recent single cell transcriptomic studies found unique expression profiles for excitatory neurons in frontal versus occipital cortical areas in mature mice (Saunders et al., 2018;Tasic et al., 2018). This suggests that excitatory neuron types are unique in these regions, and thus might form area-specific local circuits that result in functional specialization. However, despite their differences, the major classes of excitatory neurons are conserved throughout the cortex (Yao et al., 2021). In agreement, we found greater differential expression of circuit-related genes between major classes within a cortical region than between best matched classes across regions. However, our analysis of class-relevant genes within each region revealed several that may lead to region-specific circuits. Thus, whether excitatory neurons form conserved circuit motifs organized by class remains an open question. An important next step is to determine which of these genes mediates local circuits versus long-range afferent and efferent connectivity. Our analysis provides several candidate genes to investigate these possibilities in future studies.

In contrast to excitatory neurons, inhibitory interneurons are produced from progenitor pools within transient embryonic structures in the ventral telencephalon (Lavdas et al., 1999;Nery et al., 2002;Butt et al., 2005;Miyoshi et al., 2007;Miyoshi et al., 2010). Immature interneurons migrate to the dorsal telencephalon, where they transition to tangential migration to disperse across the entire developing cortex and hippocampus (Anderson et al., 1997;Ang et al., 2003;Marin, 2013;Mayer et al., 2015;Pelkey et al., 2017). Because interneurons throughout the mature cortex have a common embryonic origin, it is not surprising that the transcriptomic profiles of major classes are uniform from the rostral to caudal poles (Saunders et al., 2018;Tasic et al., 2018). Thus, interneurons have been assumed to form canonical circuit motifs that depend on their class, regardless of cortical or hippocampal region (Kepecs and Fishell, 2014;Pelkey et al., 2017;Huang and Paul, 2019). However, regional cues play a role in interneuron differentiation and maturation (Ishino et al., 2017;Quattrocolo et al., 2017;Petros, 2018), and relationships between major classes of excitatory and inhibitory neurons determine cortical circuit organization. For example, PT-type cells guide the radial migration of PV+ and SST+ interneurons (Lodato et al., 2011), and IT-type cells guide migration of VIP+ interneurons (Wester et al., 2019). Thus, it is possible that variations in major excitatory classes in different cortical regions result in unique inhibitory circuit motifs. Indeed, recent work using monosynaptic rabies tracing suggests both long-range afferent and local circuit connectivity of PV+ and SST+ interneurons is region dependent (Pouchelon et al., 2021). We found that expression of circuit-related genes depends mostly on major interneuron class rather than cortical region. Thus, our analysis supports the hypothesis that interneurons engage in canonical circuit motifs across the brain. However, we did identify genes that were unique for each interneuron class between ALM and VISp and may guide regional inhibitory circuit motifs.

### Directions for future studies of neocortical circuit organization

Our analysis provides clear targets for future investigations of cortical circuit motifs and the molecular mechanisms underlying their organization and function. Importantly, we associated genes that regulate synapse formation, maintenance, and plasticity with specific classes of excitatory and inhibitory neurons. Thus, our analysis can be used to test hypotheses regarding the function of these genes in different cell types and brain regions using Cre mouse lines that allow conditional deletion in excitatory projection neurons (Gerfen et al., 2013;Matho et al., 2021) and inhibitory interneurons (Taniguchi et al., 2011;He et al., 2016). For example, *cdh13* encodes a cadherin previously shown to be selectively expressed in PT-type excitatory neurons to guide axon targeting (Alcamo et al., 2008;Hayano et al., 2014). Importantly, our analysis identified *cdh13* as a circuit-related gene that is class-relevant to L5 PT cells in both ALM and VISp (**Fig. 7A**). Interestingly, in the hippocampus, *cdh13* is preferentially expressed in the presynaptic terminals of SST+ dendrite-targeting interneurons, where it modulates GABA release onto excitatory pyramidal neurons (Rivero et al., 2015). In the neocortex, we also identified *cdh13* as class-relevant to SST+ interneurons (**Fig. 7C**). In deep cortical layers, SST+ interneurons and PT-type excitatory neurons form a disynaptic feedback inhibition circuit motif (Le Be et al., 2007;Silberberg and Markram, 2007). An intriguing hypothesis is that *cdh13* plays an important role in the formation or maintenance of this motif.

We identified few genes that uniformly delineate excitatory or inhibitory classes across ALM and VISp (**Fig. 10**). However, the genes we did find were previously shown to play important roles in the regionalization of the rostral and caudal neocortical poles. For example, *id2* is preferentially expressed in the caudal half of cortex (Rubenstein et al., 1999), and *lmo4* is preferentially expressed in motor cortex relative to visual cortex (Cederquist et al., 2013). Interestingly, our analysis also identified *tenm2* as being preferentially expressed in VISp relative to ALM (**Fig. 10A**). Strikingly, *tenm2* knock out mice exhibit visual but not motor deficits (Young et al., 2013), which provides a functional validation of our results. Finally, we identified novels genes that may regulate cortical regionalization. These include the histone H3 variant *cenpa*, which we found is preferentially expressed in interneurons in VISp (**Fig. 10B**). CENP-A has mostly been studied for its role in defining centromeric nucleosomes and supporting chromatin compaction in preparation for kinetochore binding and chromosomal segregation during cell division (Zhou et al., 2022). However, recent work found that CENP-A, and other components of mitosis, are repurposed in neurons to regulate circuit formation (Del Castillo et al., 2019;Zhao et al., 2019). Indeed, in *Drosophila*, CENP-A mutants show disruptions in synaptic development and neurite growth (Zhao et al., 2019). The role of CENP-A in cortical circuits is unknown, but our analysis suggests it plays a specialized role in visual cortex inhibitory circuitry.

### Implications for understanding the circuit mechanisms of autism

A challenge to understanding ASD is its broad range of impairments in social-communication, cognition, and sensorimotor functions, due to disruptions in neural circuits across the brain (Marco et al., 2011;Whyatt and Craig, 2013;Donovan and Basson, 2017;Velmeshev et al., 2019;Chowdhury et al., 2021). A prominent hypothesis is that an imbalance in the ratio of excitation to inhibition (E/I) corrupts information processing within these circuits, leading to pathology (Rubenstein and Merzenich, 2003;Sohal and Rubenstein, 2019). Indeed, accumulating evidence suggests inhibitory interneurons play an important role in neurodevelopmental disorders (Goff and Goldberg, 2019;Mossner et al., 2020;Contractor et al., 2021;Goff and Goldberg, 2021;Nomura, 2021;Tang et al., 2021). However, the circuit mechanisms underlying ASD remain unclear. To develop therapies that can potentially restore E/I balance, it is crucial to identify specific circuit motifs that are vulnerable in ASD. Here, we mapped candidate ASD risk genes that encode proteins necessary for synaptic connectivity and plasticity to major neuronal classes in both ALM and VISp. Our analysis implicates cell-type specific circuit components that are conserved across the cortex or are region-specific. Thus, in some patients targeting a canonical circuit motif may alleviate multiple symptoms; however, in others, therapies may require region-specific interventions.

Interestingly, we identified many circuit-related ASD risk genes in IT-type excitatory neurons and VIP+ and SST+ inhibitory interneurons that were conserved between ALM and VISp. Thus, these are attractive candidates underlying impaired canonical circuit motifs. Importantly, enrichment in these cell types is consistent with recent single cell transcriptomic analysis in patients with ASD (Parikshak et al., 2013;Velmeshev et al., 2019). In humans, Velmeshev et al. (2019) highlighted *stx1a* in IT cells and *grik1* in VIP+ cells as key genes that are down or upregulated, respectively, in ASD patients. Our analysis identified these ASD risk genes as relevant to the same neuronal classes in mice. Thus, our work is consistent with findings in ASD patients and has great potential to guide future translational studies.

### Limitations of the study

We inferred molecular mechanisms of synaptic connectivity between major neuronal classes based on relative levels of mRNA expression. However, this approach suffers from some limitations (discussed in detail in Sudhof (2018)). First, mRNA and protein expression levels do not always correlate. Thus, some circuit-related genes identified in our differential expression analysis may not be relevant to connectivity biases among specific neuronal classes. Second, single-cell sequencing can have a shallow read depth and fail to detect alternative splicing events. Thus, subtle yet important gene expression differences between neuronal classes are likely not represented in the data we analyzed. However, the purpose of our analysis was to provide strong candidate molecular mechanisms to be verified and investigated in future studies. Our stringent approach to assign each gene to a distinct neuronal class likely identified many that play specialized roles in circuit organization.

Our strategy filtered out several genes for which individual tests were statistically significant (**Fig. 5**). Some of these are undoubtedly important for establishing circuit motifs. An example is *cxcl12*, which encodes a cytokine necessary for attracting the axon terminals of PV+ interneurons (Wu et al., 2016). Selective expression of *cxcl12* by PT cells, but not IT cells, results in biased inhibition from PV+ interneurons (Lee et al., 2014;Wu et al., 2016). Importantly, our analysis found significantly greater expression of *cxcl12* in L5 PT than L5 IT classes (not shown). However, this gene was not differentially expressed in other tests between classes, and thus was not defined as “class-relevant”. Although our analysis filtered this gene, we emphasize that our stringent criteria highlight many others that are strong candidates for future investigation.

Finally, each major neuronal class contains unique subtypes that may form specialized microcircuit motifs (Tasic et al., 2018;Chen et al., 2019;Kim et al., 2020;Cheung et al., 2021;Que et al., 2021). In future studies, it will be interesting to use the metadata provided by the Allen Institute to further refine our analysis to consider these subtypes.

## Conclusions

Our analysis leveraged the Allen Institute’s single-cell transcriptomic datasets to reveal candidate genes underlying cell-type-specific circuits across functionally distinct cortical regions. We conclude that major neuronal classes likely establish many canonical circuit motifs that are conserved across the neocortex. However, we also identified many genes that may regulate region-specific motifs in a cell-type-specific manner. Our analysis will hopefully serve as a resource for future studies of cortical circuit connectivity and the molecular mechanisms underlying its organization.

## METHODS

### Datasets

We downloaded the Allen’s ALM and V1 – SMART-seq (2018) datasets (https://portal.brain-map.org/atlases-and-data/rnaseq). Using the scRNA-seq toolkit Seurat (Stuart et al., 2019) and provided metadata, we filtered for glutamatergic and GABAergic neurons from each brain region to create four separate Seurat objects. We merged glutamatergic and GABAergic objects from separate regions to create two final objects for analysis. We followed the Seurat standard pre-processing workflow for QC, normalization, identification of highly variable features, scaling, dimensionality reduction, and UMAP clustering.

### Conducting differential expression tests

Using brain region and class metadata, expression profiles of cells from each brain region (ALM or VISp) and selected major classes (L2/3 IT, L5 IT, L5 PT, and L6 CT for glutamatergic object; Pvalb, Sst, and Vip for GABAergic object) were compared in a pairwise manner (**Figs. 2A, B**). Comparisons were performed using the Seurat function FindMarkers which implements a non-parametric Wilcoxon rank-sum test for differential expression. For comparisons between different classes within a brain region (**Fig. 2A**), we subsetted each object for cells in either ALM or VISp before performing FindMarkers. For example, to compare SST and PV cells in ALM, we first subset the GABAergic object for cells in ALM and used the class metadata to perform the test. For comparisons between the same classes across brain regions (**Fig. 2B**), we subset the glutamatergic or GABAergic object for the class of interest before performing FindMarkers. For example, to compare SST cells across ALM and VISp, we subset the GABAergic object for SST cells and used the region metadata to perform the test. Equivalent comparisons between different classes within ALM or VISp were conducted in the same order (e.g. GABA ALM SST vs PV is equivalent to GABA VISp SST vs PV). This allows the log fold change values of differentially expressed genes to be directly compared for equivalent tests in ALM and VISp. Differentially expressed genes were selected with an adjusted P-value cut-off of <0.05 based on Bonferroni correction using all genes in the dataset.

### Approach for normalizing differential expression counts

To determine the maximum number of genes that could be differentially expressed in each pairwise comparison, we used the AverageExpression function in Seurat. This function returns the averaged expression of each gene across all cells in an object (log normalized expression values were exponentiated before averaging). For comparisons between different classes within a brain region (**Fig. 2A**), we subsetted the glutamatergic or GABAergic object for a pair of cell-types before performing AverageExpression. For example, to determine the number of unique genes expressed by SST and PV cells, we first subset the GABAergic object for both classes and used the region metadata to calculate average expression in ALM and VISp. For comparisons between the same class across brain regions (**Fig. 2B**), we used the class metadata to identify cells in the glutamatergic or GABAergic object before performing AverageExpression. For example, to determine the number of unique genes expressed by SST across ALM and VISp, we labeled cells in the GABAergic object by class and used the averaged expression results pertaining to SST cells. For each comparison, we removed genes with an average expression value of 0. This enabled us to determine the total number of unique genes that are expressed by each pair of cell-types in **Figures 2A, B**. We term these genes “union” since they are the union of the set of genes that are expressed by a pair of cell-types. Using the differential expression and union data for each pairwise test, we calculated the proportion of genes that are differentially expressed (e.g. out of 33581 unique genes expressed by L2/3 IT and L5 IT cells in ALM, 1.4% (465 genes) are differentially expressed). We organize these results in a matrix presented in the Circos plot in **Figure 2C** (Krzywinski et al., 2009).

### Filtering for circuit-related genes

To identify genes that are circuit-related, we used the classification software PANTHER 16.0 (Mi et al., 2021). We uploaded differentially expressed and union genes, selected Mus musculus as the reference organism, and ran a statistical overrepresentation test (GO biological process complete) with no corrections. Genes labeled under the gene ontology categories “Cell-cell adhesion”, “Regulation of cell-cell adhesion”, and “Regulation of trans-synaptic signaling” were selected. We annotated and filtered for selected genes in the differential expression and union gene sets. For every GO category and pairwise test, we normalized the number of differentially expressed genes to the number of GO union genes in the manner described above. We present these results in the Circos plots in **Figure 3** (Krzywinski et al., 2009).

### Labeling for clinically relevant genes

We downloaded the 2021 list of autism risk genes from the SFARI Human gene module and labeled differentially expressed genes that are risk genes. Risk genes from all categories (“Suggestive Evidence”, “Strong Candidate”, “High Confidence”, and “Syndromic”) were considered.

### Strategy for determining the regional-specificity of each gene

For our analysis, we consider a differential expression datapoint as a coordinate of the following features: Comparison (Glut/GABA ALM/VISp Class 1 vs Class 2 or Glut/GABA ALM vs VISp Class 1), gene, log fold change (logFC) value, relevant GO terms, and relevant conditions. We compiled results for glutamatergic and GABAergic differential expression data that were filtered for circuit-related genes. We created the tool DE_Collapser in Python to identify differential expression data that is “ALM-specific”, “VISp-specific”, “conserved” or “unique” (**examples in Fig 4A**). This was accomplished using the following algorithm: Analyzing the compiled differential expression data, DE_Collapser iterates through pairs of comparisons to identify comparisons that are equivalent across brain regions (e.g. GABA ALM SST vs PV is equivalent to GABA VISp SST vs PV). It then subsets for their corresponding differential expression data and identifies the intersection of the sets of genes that are differentially expressed in each comparison. For each of these genes, DE_Collapser determines if the relative expression between cell-types is either consistent across comparisons or different by using the sign of the logFC. The logFC values can be used because classes were compared in the same order in equivalent differential expression tests. Genes exhibiting a consistent relationship across regions are labeled ‘conserved’. For example, *shisa6* is enriched in VIP cells relative to PV cells in ALM and VISp (**Fig. 4Aiii**). This is indicated by a positive logFC value in the ALM and VISp comparisons. Genes exhibiting a relationship that is reversed across regions are labeled ‘unique’. For example, L5 IT cells have greater expression of *calb1* relative to L5 PT cells in ALM but this relationship is reversed in VISp (**Fig. 4Aiv**). This is indicated by opposite signs of the logFC values for the comparison in ALM versus that in VISp. For conserved and unique genes, DE_Collapser computes the average of the logFC values from equivalent comparisons and converts a pair of differential expression datapoints for that gene into one datapoint with the averaged logFC value. Datapoints for unique genes are also separated into a distinct subset that preserves raw logFC values for comparisons in each region. For genes that are differentially expressed in only one of two equivalent comparisons, DE_Collapser identifies if they are differentially expressed in the ALM comparison (label ‘ALM-specific’; **Fig. 4Ai**) or VISp comparison (label ‘VISp-specific’; **Fig. 4Aii**). Examples of applying DE_Collapser on raw datapoints is represented by the set of arrows between **Figure 5A** and **Figure 5B**. The output of DE_Collapser for each pair of comparisons is compiled into a new dataset combining conserved and unique data with averaged logFC values and ALM- or VISp-specific data with raw logFC values. Raw logFC values for ‘Unique’ genes remain accessible in a separate output. We calculated the total number of conserved, unique, and region-specific genes in each comparison **(Figure 4B**).

### Strategy for class-relevance analysis

All of the following class analyses presented in **Figures 5-10** were performed separately for glutamatergic and GABAergic data. To determine if differentially expressed genes map to class features, we created Subclass Identifier (SCID). This algorithm takes the output of DE_Collapser including conserved or region-specific data. It separates comparisons into the order of their constituent classes (e.g., VIP vs PV – First class = VIP; Second class = PV). Based on the logFC values, SCID identifies which class has greater or lesser relative expression of each gene: If the logFC > 0 for gene A in a VIP vs PV comparison, then the first class (VIP) has greater expression and second class (PV) has lesser expression. If the logFC < 0 for gene B in a VIP vs PV comparison, then the second class (PV) has greater expression and first class (VIP) has lesser expression.

To assist with classification, we define sets for the minimum comparisons that a gene needs to be differentially expressed in to be in consideration for mapping to a class: L2/3 IT = {‘L2/3 IT vs L5 IT’, ‘L2/3 IT vs L6 CT’, ‘L2/3 IT vs L5 PT’}

L5 IT = {‘L2/3 IT vs L5 IT’,’L5 IT vs L5 PT’,’L5 IT vs L6 CT’}

IT = {‘L2/3 IT vs L5 PT’, ‘L2/3 IT vs L6 CT’, ‘L5 IT vs L5 PT’,’L5 IT vs L6 CT’}

L5 PT = {‘L2/3 IT vs L5 PT’,’L5 IT vs L5 PT’,’L5 PT vs L6 CT’}

L6 CT = {‘L2/3 IT vs L6 CT’,’L5 IT vs L6 CT’,’L5 PT vs L6 CT’}

VIP = {‘VIP vs PV’, ‘VIP vs SST’}

SST = {‘SST vs PV’, ‘VIP vs SST’}

PV = {‘SST vs PV’, ‘VIP vs PV’}

The following steps comprise the key computations of the algorithm which we demonstrate with an example considering a hypothetical gene X. We refer to a comparison set as all of the comparisons that a gene appears differentially expressed in.

1. SCID iterates through the differential expression data for each gene and determines the list of classes that differentially express the gene. If L2/3 IT and L5 IT are identified, IT is added to the class list. Example: If gene X is differentially expressed in the comparison set {‘VIP vs PV’, ‘VIP vs SST’, ‘SST vs PV’}, then SCID identifies SST, VIP, and PV as the class list.
2. For each class in the list, SCID iterates through pertinent class sets that we define above. It identifies class sets that are a subset of the comparison set associated with the gene. Example: For gene X, the VIP, {‘VIP vs PV’, ‘VIP vs SST’}; SST, {‘SST vs PV’, ‘VIP vs SST’}; and PV {‘SST vs PV’, ‘VIP vs PV’} class sets would each be identified as a subset of the comparison set {‘VIP vs PV’, ‘VIP vs SST’, ‘SST vs PV’}.
3. For each class set, SCID filters the differential expression data for comparisons that are included in the set. Example: For the VIP set, SCID filters for Gene X data involving ‘VIP vs PV’, ‘VIP vs SST’ comparisons; for the SST set, it filters for Gene X data involving ‘SST vs PV’, ‘VIP vs SST’ comparisons; and for the PV set, it filters for Gene X data involving ‘SST vs PV’, ‘VIP vs PV’ comparisons.
4. In each filtered dataset, SCID evaluates the information on which class has greater or lesser expression of a gene in each comparison. This enables it to determine if the class (e.g. VIP) corresponding to the class set (e.g. {VIP vs PV’, ‘VIP vs SST’}) has consistent expression relative to all other classes (e.g. PV and SST). We refer to these as ‘class-relevant’ genes. We label class-relevant genes for their class features and the corresponding expression relative to all other classes for each feature **(Figs. 5C, 7-9**). Genes that are not class-relevant remain unlabeled. Example: If gene X is expressed greater in VIP cells relative to PV and SST cells, and expressed less in SST cells relative to VIP and PV cells, gene X is labeled accordingly: up in VIP cells and down in SST cells.

Note: Given the low number of genes in the unique subset, we searched for class-relevant genes directly. No genes were identified. Also, per the defined class lists and differential expression analysis, SCID identified features are relative to either glutamatergic or GABAergic classes.

### Strategy to identify class-relevant genes that involve conserved and region-specific data

To account for class features that may be overlooked if only considering region-specific data (**Fig. 6**), we created CortexSCID. This algorithm considers the results of performing SCID on both the conserved data and the raw ALM or VISp data. The latter contains all of the ALM or VISp data not just that which is region-specific. CortexSCID identifies genes that are present in the conserved set and ALM or VISp sets and removes genes that have the same comparison sets and class features. The remaining genes have one or more class features that are distinct to ALM or VISp. We refer to these as * genes. CortexSCID identifies * genes, their class features, and corresponding relative expression. It updates the ALM or VISp differential expression data by preserving only genes and their class relevance that are region-specific. * genes are only identified if the same gene is relevant to different classes in a conserved and region-specific manner (e.g. *egr1* in **Fig. 6**).

### Identifying genes that are biased to ALM or VISp

Using the differential expression data for comparisons between the same class across ALM and VISp **(Fig. 2B**), we queried for genes that are enriched across multiple glutamatergic or GABAergic classes in ALM relative to VISp or vice versa **(Fig. 10**). For example, we identified *cenpa* as being enriched in VIP, PV, and SST cells in VISp relative to their equivalents in ALM (**Fig. 10B left, Fig. 10D**). We also searched for genes that are differentially expressed by VIP cells but not PV or SST cells across brain regions (**Fig. 10B right, Fig. 10 E**) or PV and SST cells but not VIP cells across brain regions (no such genes identified). The Violin plots for *tenm2, cenpa*, and *igf1* were created in Seurat (**Fig. 10C-E**).

### Heatmap creation

We created the function Morpheus Prepper (MorphPrep) to streamline class and clinical analyses and organize results as matrices for heatmap visualization in Morpheus (https://software.broadinstitute.org/morpheus).

